# Activation of Macrophages by CpG DNA and LPS: an FTIR Spectroscopic Study

**DOI:** 10.1101/2020.03.12.987354

**Authors:** Emrulla Spahiu, Senol Dogan, Jörg Schnauß, Mayda Gursel, Feride Severcan

**Affiliations:** Department of Biological Sciences, Middle East Technical University, 06800, Ankara, Turkey; Leipzig University, Faculty of Physics and Earth Sciences, Peter Debye Institute for Soft Matter Physics, Linnéstraße 5, 04103 Leipzig, Germany; Fraunhofer Institute for Cell Therapy and Immunology, 04103 Leipzig, Germany; Department of Biophysics, Faculty of Medicine, Altinbas University, Istanbul, Turkey

**Keywords:** Ftir, infrared, spectroscopy, innate immunity, raw 264.7, activation, macrophages, cpg dna, lps

## Abstract

The innate immune response triggered by CpG DNA can improve host survival following pathogen challenge. Whether CpG ODN-mediated immune activation leads to global molecular changes in cells that are detectable by FTIR spectroscopy is currently unknown. Here, we used Attenuated Total Reflectance (ATR) Fourier Transform Infrared (FTIR) spectroscopy to monitor the molecular changes in murine macrophage RAW 264.7 cells upon activation with CpG DNA and lipopolysaccharide (LPS). By PCA analysis, we identified the sources of variation to follow with detailed spectral analysis. CpG DNA and LPS treatment increase the total nucleic acid concentration from the early periods post-activation, and DNA synthesis follows RNA synthesis. RNA-specific peak shows the activation state of macrophages in early periods post-treatment. CpG DNA and LPS result in an initial rapid increase in the total protein concentration, leveling off two hours post-activation. Both activated groups increase the concentration of fatty acids, triglycerides, and cholesterol, pointing out to a shared synthesis pathway and *de novo* lipogenesis. This study, for the first time, demonstrates the use of FTIR spectroscopy as an independent modality to monitor the activation dynamics of murine macrophages upon activation with CpG DNA and LPS.

## 1. Introduction

FTIR spectroscopy has developed into a powerful biomedical tool by allowing non-destructive, rapid, and high-throughput analysis of biological systems in both qualitative and quantitative manners [1]. From a single spectrum, all delicate alterations in both the conformation and concentration of vital macromolecules, such as nucleic acids, proteins, and lipids, can be revealed. Thus, the infrared (IR) spectrum serves as a “fingerprint” for all metabolites of the cell system. For an IR spectral peak, the area under the curve provides quantitative (concentration) data, while the specific absorption frequencies reveal the nature of the chemical bond and molecular environment [2]. The absorbance of IR light by water molecules, however, is a limitation of infrared spectroscopy since it superimposes other signals of the aqueous cell systems [3]. The development of the Fourier transform procedure overcame this disadvantage. ATR unit offers a handy use of spectrometer since the sample is directly put onto the ATR crystal; no sample preparation step is required. In addition, a very little amount of sample is enough to obtain a good quality spectrum [4, 5].

ATR-FTIR spectroscopy has multiple applications in hematology and was used in the analysis of live cell activation [6, 7], the screening of numerous diseases [8–11], the response of cells to chemical stress [12], the identification of disease states in the early stages [13–15] and cellular differentiation [16].

The macrophage cells used in this study constitute the front line of the immune system defending the organism against invaders. They have a high structural and functional plasticity that allows them to continually inspect the body and respond to challenges in it [17–19]. An intriguing challenge is bacterial DNA containing unmethylated CpG oligodeoxynucleotides (CpG DNA) [20, 21]. CpG DNA triggers the proliferation of macrophages, the production of pro-inflammatory cytokines and chemokines, the induction of Th1 reactions, the repression of Th2 reactions, the activation of antigen-presenting cells and the proliferation of B cells [22].

The same response is mimicked by synthetic CpG oligodeoxynucleotides (ODN) containing CpG motifs, serving as potential immune adjuvants with therapeutic uses [23]. The innate immune response triggered by CpG containing ODNs can improve host survival following pathogen challenge. However, how CpG ODN-mediated immune activation impacts relative amounts of nucleic acids, lipids, proteins, and other metabolites has not been studied.

Lipopolysaccharide (LPS), on the other hand, is one of the main constituents of the cell wall of gram-negative bacteria [24]. It occupies approximately 75% of their external surface, rendering it a good indicator of bacterial infection in the body [25]. LPS stimulates the monocytes and macrophages, which respond by releasing a wide variety of biological response mediators, including platelet-activating factor, enzymes, free-radicals, and a mix of pro-inflammatory cytokines, namely tumor necrosis factor-a (TNF-a) and interleukins (IL-1, IL-6, IL-8, and IL-12) [26]. As a result of these responses, microbial infection is prevented or delayed [27]. The recognition process of LPS is carried out through the TLR4 pathway sharing high homology with TLR-9 signaling carried out by CpG DNA [28]. Both need MyD88 as the adaptor protein and activate NF-kB as well as AP-1 transcription factors [29].

Understanding the rules governing cellular responses to stimulatory motifs can foster the design of ODNs for therapeutic uses [30]. Even though the trending “omics” techniques yield precise quantitative data on specific molecules rather than the bulk analysis done by FTIR spectroscopy, they require complex sample preparation steps that can result in alterations of chemical ratios and post-translational modifications in lipidomics [31, 32] and proteomics [33, 34]. The simple, rapid, and non-invasive experimental setup using bulk FTIR analysis proves to give a broad view of the relative amounts of nucleic acids, lipids, proteins, and other metabolites that are not readily accessible by other techniques [35, 36].

In the present study, ATR-FTIR spectroscopy was used to examine the global molecular changes in murine macrophage cells upon treatment with CpG DNA and LPS. Using this technique, many cells were probed simultaneously, yielding statistically significant data sets.

## 2. Results and Discussion

A comprehensive spectrum of resting control RAW 264.7 cells together with its second derivative in 4000-650 cm^−1^ IR spectral region is presented in Figure A1. FTIR spectrum derived from murine macrophage cells, points to a complex composition, containing peaks from lipids in the 3000-2800 cm^−1^ region, proteins (1700–1600 cm^−1^), nucleic acids (1238 cm^−1^, 1110 cm^−1^, and 1085 cm^−1^) and numerous other metabolites. Table A1 shows the peak assignments based on literature, and Table A2 shows the measurements of area integration regarding peaks of interest (for all study groups see Table 1).

**Table 1.**
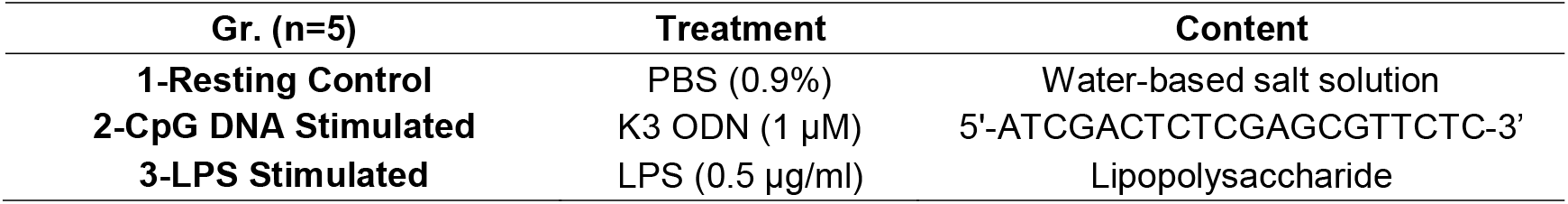
Characteristics of treatments in study groups

### 2.1. Principle Component Analysis (PCA) Resulted in Good Approximation of the Experimental Results

PCA is used to reduce the dimensionality of large data sets, thereby drawing logical conclusions on the significant changes between the study groups. It is an unsupervised multivariate data analysis (MDA) method, working without initial information input regarding the datasets [37]. The two useful outputs of PCA analysis, Scores Plot—showing the clusters of similar data—and Loadings Plot—showing the sources of highest variation (in our case, the wavenumbers), gave useful information for the initial steps of data analysis [38]. PCA in our samples was conducted on the data matrix obtained from spectral readings of 0.5 cm^−1^ resolution. Principal Component (PC)-1 gave a variability of 83% and provided information for the vast source of variation (Figure 1.A) when compared to PC-2, which had a variability of 11%. Loadings Plots (Figure 1.B and 1.C) show the 1750-850 cm^−1^ IR region having numerous peaks with high variation among samples. The patterns observed in the Scores Plot (Figure 1.A), on the other hand, show a considerable amount of samples being clustered according to the treatment that they obtained in two PC directions. PCA results — apart from validating the spectral data — guided us in selecting the peaks for detailed peak analysis.

**Figure 1.**
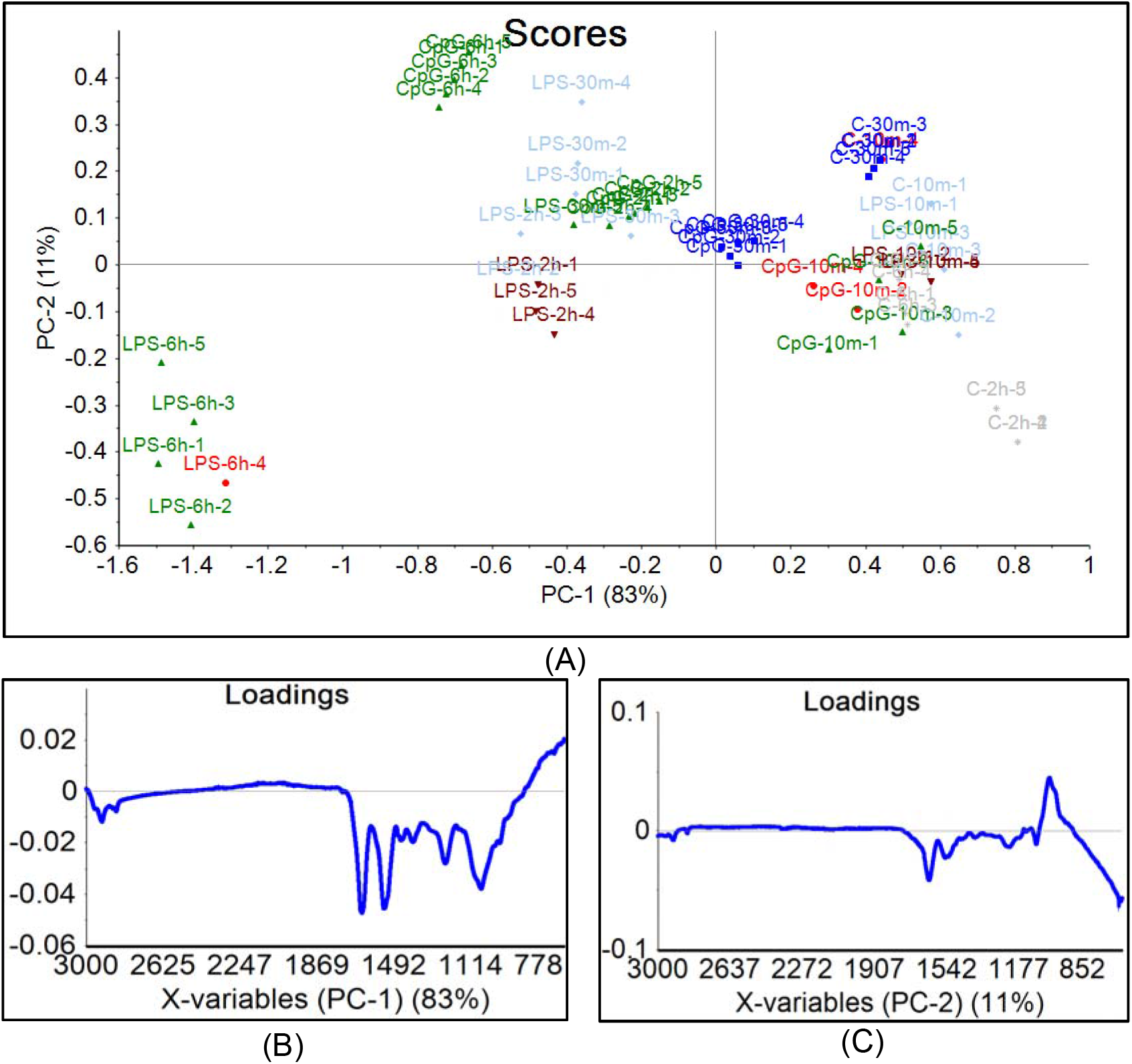
The initial evaluation of results was carried out by PCA analysis. PCA Scores plot of FTIR spectra in the 3000-650 cm^−1^ IR region, shows a considerable amount of samples being clustered according to the treatment that they obtained. Control group (C), CpG-treated group (CpG), Lipopolysaccharide-treated group (LPS). Sample designation: Treatment-time of treatment-number of replicate. (**A**). Loading plots of Principal Component (PC)-1 and PC-2 point out to many peaks in 1750-850 cm^−1^ region containing high wavenumber variations. PC-1 has a variability score of 83% (**B**), and PC-2 has a variability of 11% (**C**).

### 2.2. CpG DNA Induces RNA and DNA Synthesis During Early Stages of Macrophage Activation

From the spectrum 10 min after the treatment (Figure 2.A), activation-induced changes can be detected. The peaks related to nucleic acids—phosphate vibration (1238 cm^−1^), and symmetric PO^−2^ stretching (1085 cm^−1^)—show prominent differences between the control and stimulated groups. The peak integration values (corresponding to concentration) are shown as bar diagrams in Figure 2.B. CpG DNA stimulation induced significant changes in nucleic acid content (1238 cm^−1^ peak integration) of macrophages starting from 10 min until 6 h after treatment (p < 0.1*). It is well-known that a potential adaption of DNA synthesis following treatment becomes prominent only after several hours post-stimulation [39]. Thus, increases in nucleic acid content at early stages can be attributed to an increase in RNA content. This is verified by using the RNA-specific peak (ribose vibration), appearing at 915 cm^−1^. As depicted in Figure 2.C, stimulation with CpG DNA has significantly increased this peak value from the first minutes (p < 0.001***). This increase is maintained until 2 h after stimulation (p < 0.1*). All put together, contribution to nucleic acid-related peaks comes from DNA or RNA in a time-dependent manner. The time until 2 h after stimulation seems to be the period of extensive RNA synthesis, and as this ceases, DNA synthesis starts from 6 h after stimulation with CpG DNA. LPS, on the other hand, has an increased concentration of total RNA up to 2 h post-stimulation. In a gene expression study by Gao et.al. LPS is shown to induce as many as 263 genes during the stimulation process and that it is a potent pro-inflammatory and anti-proliferative agent [40]. The repertoire of genes being repressed covers the ones related to cell cycle, entirely consistent with previous studies where LPS was shown to inhibit DNA synthesis and elicit cell cycle arrest in macrophages [41, 42]. This validates the reason for no significant changes occurring in nucleic acid content after 6 h of stimulation and significant changes in RNA synthesis until 2 h after stimulation. In another study of murine macrophage activation using LPS [43], it was achieved to define the activation state of the cells as early as 41 min after stimulation. The results of our study deduce that by using the RNA-specific peak (915 cm^−1^), one can define the activation state of macrophages at the very first minutes of activation, filling the gap in the big picture of their activation.

**Figure 2.**
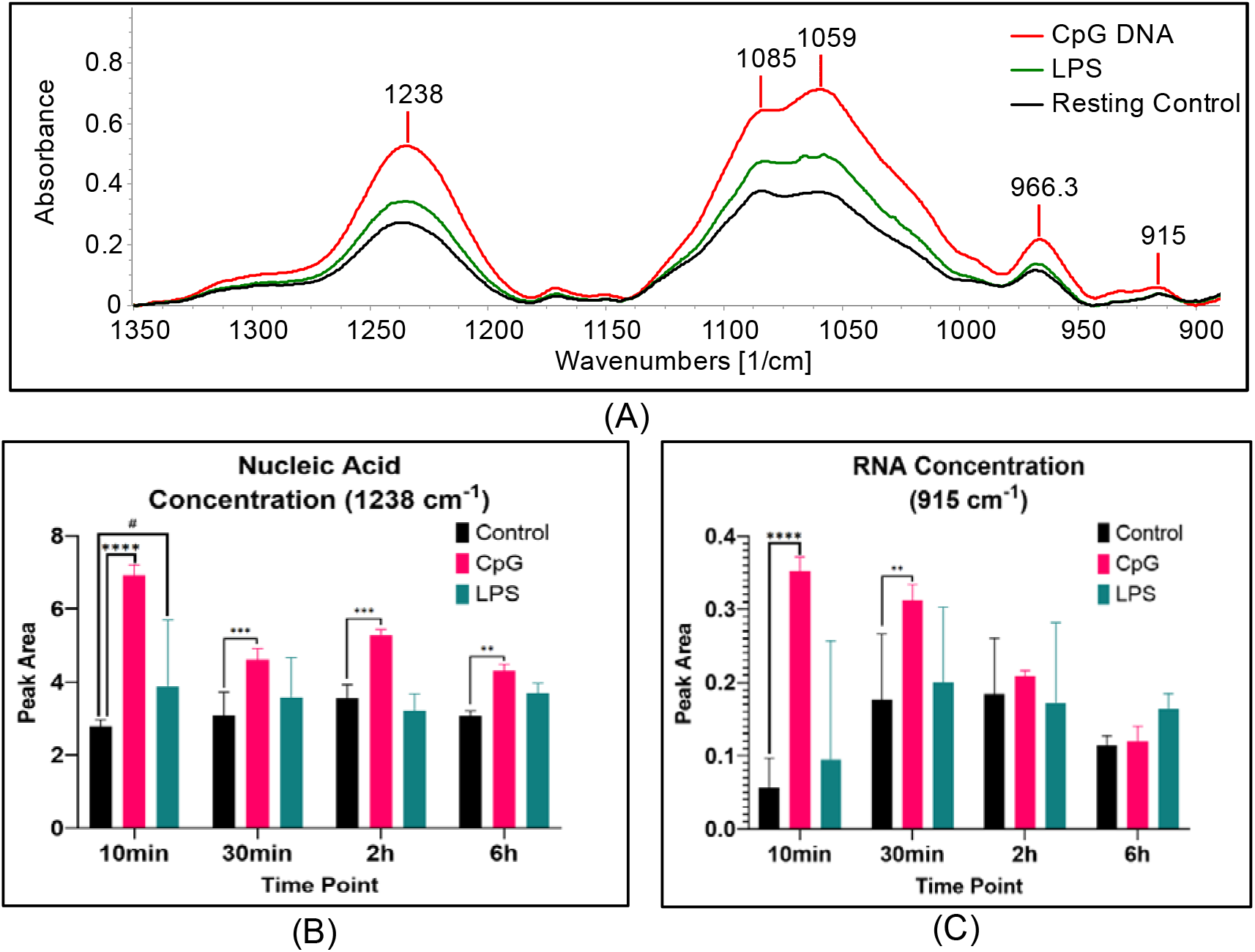
(**A**) The representative FTIR spectra showing the differences between resting (control), and activated (CpG DNA and LPS) murine macrophage cells in 1350-900 cm^−1^ spectral region, 10 min post-treatment (**B**) The integration of 1238 cm^−1^ peak (representing the phosphate vibrations in nucleic acids), shows an initial increase of nucleic acid concentration by stimulated groups (**C**) RNA concentration analysis of 915 cm^−1^ peak (ribose vibrations), shows an increased concentration of RNA (revealing the activation time) in macrophages, induced by CpG DNA and LPS. Statistical significance annotations: p ≤ 0.05 *, p ≤ 0.01 **, p ≤ 0.001 *** and p ≤ 0.0001 ****

### 2.3. CpG DNA and LPS Increase the Total Protein Concentration 2h Post-Activation, Followed by a Decrease

Amide I (1639 cm^−1^) and amide II (1540 cm^−1^) peaks are used for protein content analysis [44–47]. There is a visible alteration in the FTIR spectra of our samples in this region (Figure 3.A and 3.B). The data regarding the integration of the amide I band (Figure 3.C) further shows that there is a significant increase in the total protein content of CpG DNA treated cells from 10 min post-treatment until 2 h (p < 0.001***). There is a similar change upon LPS stimulation. The initial increase in protein concentration is reduced, starting from 2 h time point (Figure 3.C). Lines of evidence supporting this finding come from a study employing bioinformatic network analysis of microarray data [48]. It was found that major inducers (IL1A, IL1B, TNF, and IFNG) worked in a synchronized way to regulate initial gene expression; that the CpG DNA induced immune activation was short-lived; and that dynamic suppression by suppressors (IL10, MYC, NFKBIA, SOCS1, SOCS3, IL1RN, and FOS) together with a loss of induction, resulted in a decline of activation after 3 h of treatment.

**Figure 3.**
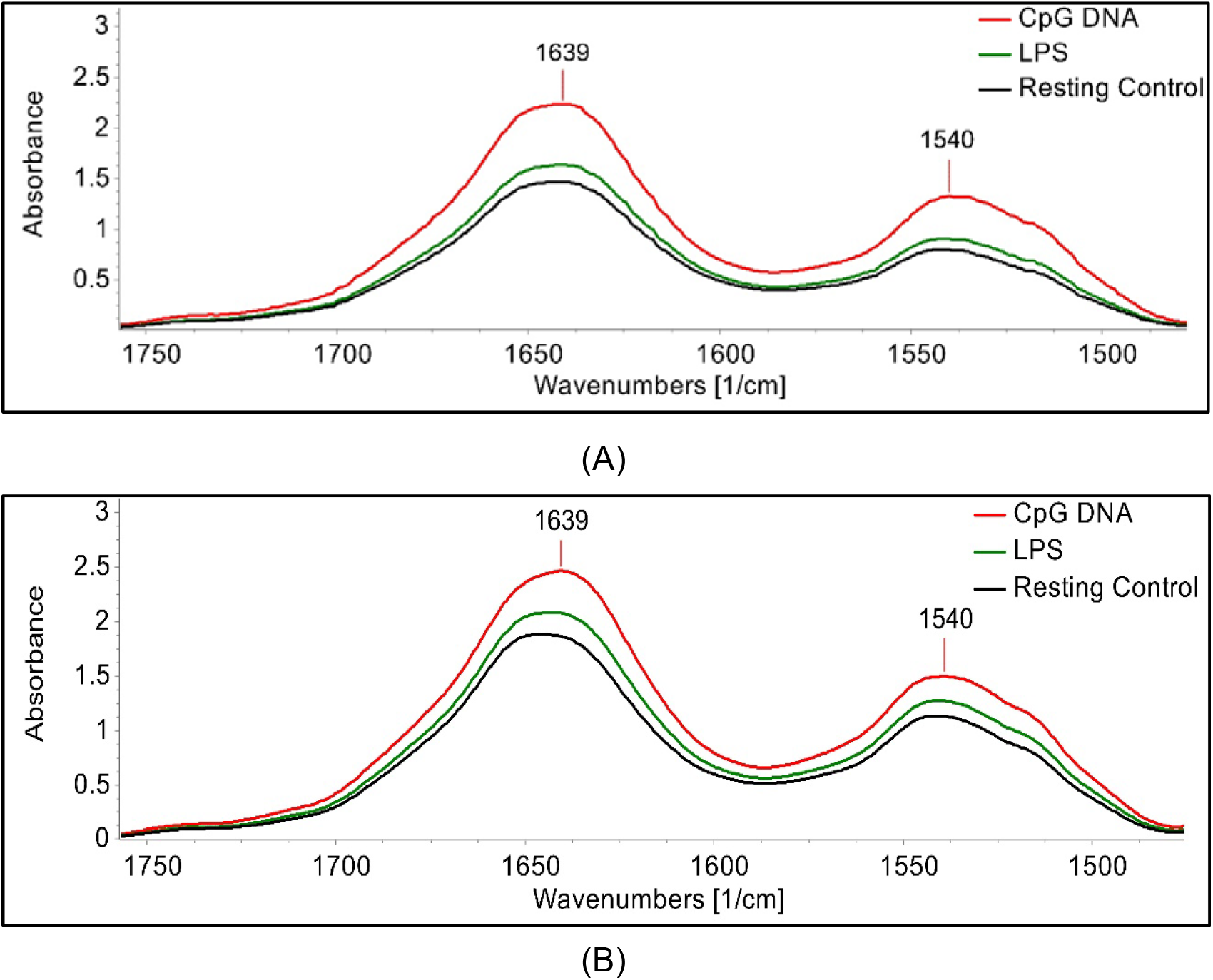
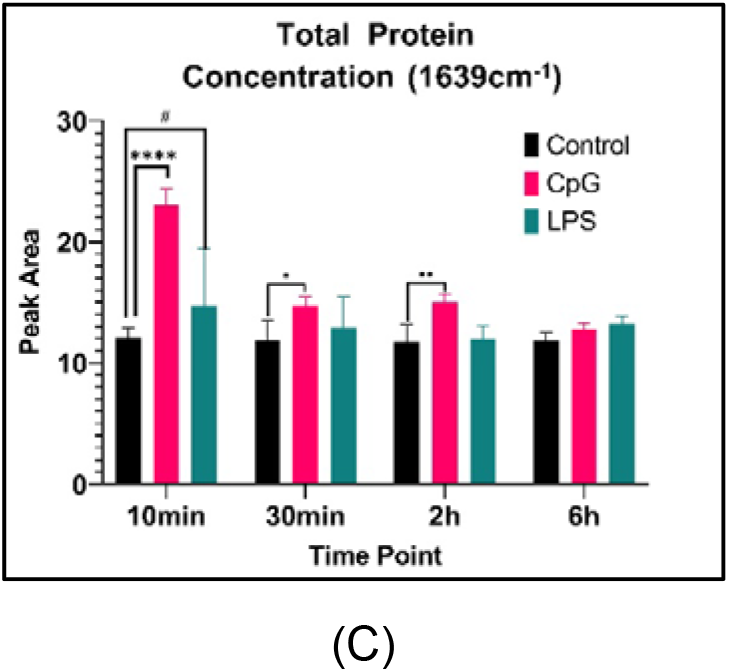
The representative FTIR spectra showing the differences between the resting control, and activated (CpG DNA and LPS) murine macrophage cells in 1750-1460 cm^−1^ spectral region (**A**) 10 min post-treatment (**B**) 6 h post-treatment (**C**) Protein concentration analysis performed by 1639 cm-1 peak integration, depicting the increase in total protein content of activated groups. Statistical significance annotations: p ≤ 0.05 *, p ≤ 0.01 **, p ≤ 0.001 *** and p ≤ 0.0001 ****.

Previously, several pathways were shown to lower the protein content during the LPS challenge. Histones were shown to be released into the extracellular medium [49], a general protein secretion via vesicles and exosomes was documented together with protein shedding from the plasma membrane [50], and the association of LPS with receptors driving the production of ROS causing high oxidative stress and degradation of oxidized proteins [51]. While these pathways contribute at a certain unknown degree to the decrease of total protein content upon LPS stimulation, our work provides an aerial view of the total protein fate and points out to the need for a study of mechanisms in CpG DNA induced protein loss in the future.

### 2.4. CpG DNA Increases the Concentration of Fatty Acids, Triglycerides, and Cholesterol

The lipid metabolism modifications based on the cell activation state [52] can also be investigated in our samples by FTIR spectroscopy. For the effect of activation in triglycerides and cholesterol, we focused on the ester C=O stretching vibrations represented in the IR spectrum at the peak near 1741 cm^−1^. There was a very significant increase in the peak during activation correlating to an increased concentration of triglycerides and cholesterol starting from the very first minutes of stimulation (Figure 4.D and 4.E). LPS stimulation, on the other hand, elevated the triglycerides and cholesterol concentration after 10 min. Cholesterol is essential for the structure (e.g., membrane fluidity, lipid rafts) and function (e.g., signaling, migration) of membranes, which have a prominent role during the activation of macrophages [53].

**Figure 4.**
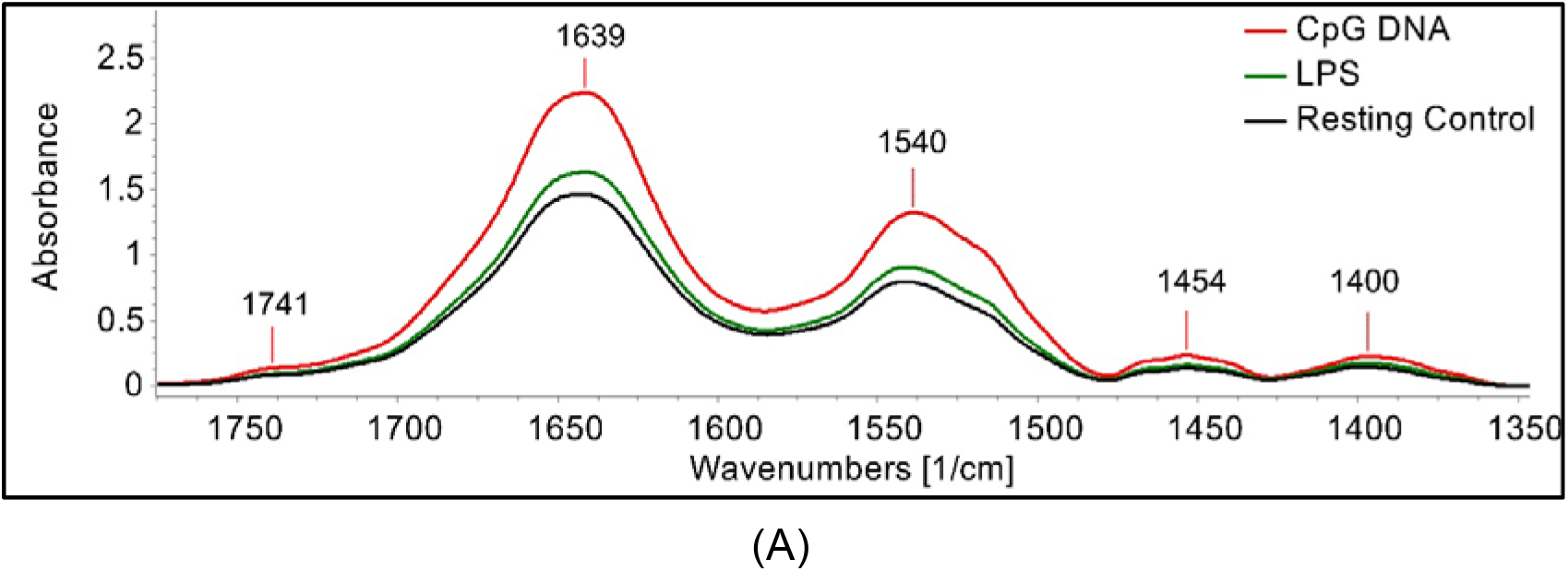
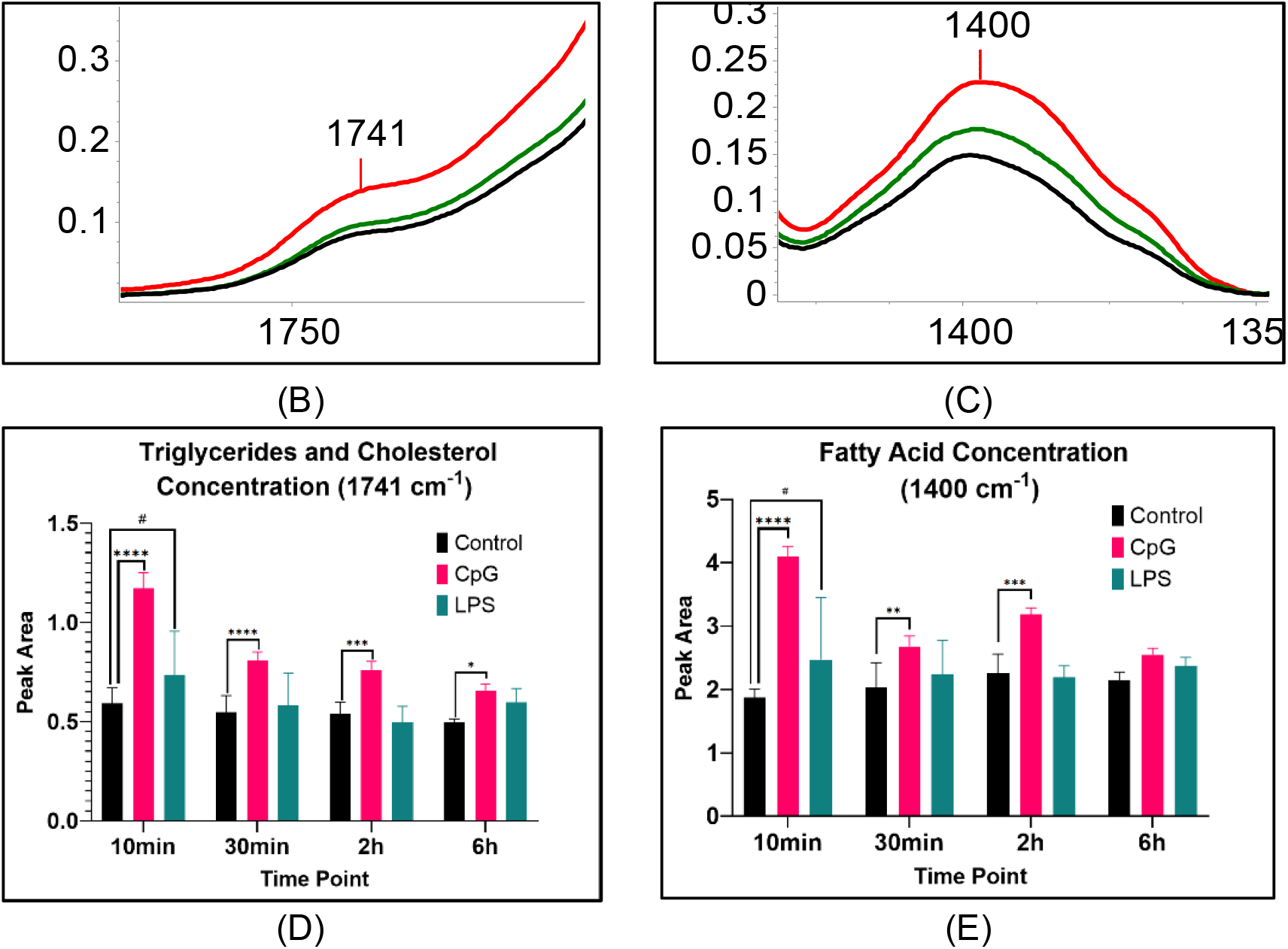
(**A**) The representative FTIR spectra of 10 min time point, showing the spectral differences between resting (control) and activated (CpG DNA and LPS) murine macrophage cells in 1750-1350 cm^−1^ region (**B**) An enlarged view of 1741 cm^−1^ peak representing ester C=O stretching vibrations of triglycerides and cholesterol esters. (**C**) An enlarged view of 1400 cm^−1^ peak, representing COO– symmetric stretching of fatty acids (**D**) Triglyceride and cholesterol concentration analysis by 1741 cm^−1^ peak integration, indicate increased concentration of triglycerides and cholesterol in activated groups (**E**) Fatty acid concentration analysis by 1400 cm^−1^ peak integration, indicate increased fatty acid concentration in activated groups. Statistical significance annotations: p ≤ 0.05 *, p≤ 0.01 **, p ≤ 0.001 *** and p ≤ 0.0001 ****.

The second dataset is derived from the COO- the symmetric stretching peak of fatty acid side chains detected in 1400 cm^−1^ (Figure 4.B and 4.C). A significant increase is observed in fatty acid concentration as well, pointing out to the previously documented *de novo* lipogenesis of pro-inflammatory reactions in macrophages [54, 55]. This previously unknown link was elaborated in a recent study on macrophage activation using different TLR agonists, where evidence was provided for a shared metabolite (Acetoacetyl-CoA) between fatty acid and cholesterol synthesis pathways [53]. Lately, CpG-induced macrophage activation was shown to trigger lipogenesis to modulate the secretory activity carried out by Endoplasmic Reticulum (ER) and Golgi. The same study drew attention to the importance of the pathway in cancer cells, where CpG-activated anti-tumor activity of macrophages is essential [56, 57].

## 3. Materials and Methods

### 3.1. Cell Preparation and Treatment

Murine macrophage-like RAW 264.7 cells were cultured at 37 °C with 5% CO_2_ in Dulbecco’s modified Eagle’s medium (DMEM; Biochrom, Germany), supplemented with 10% fetal bovine serum (FBS; Biochrom, Germany) and 2 mM L-glutamine (Biochrom, Germany). To maintain the logarithmic growth of cells, sub-culturing was done in 1/10 ratio upon reaching the acceptable confluency, generally every 2-3 days. Cells were harvested, spun at 1000 rpm 208 g for 10 min, and resuspended at a concentration of 2×10^6^/mL. Control group was treated with phosphate-buffered saline (0.9% PBS; Sigma, USA), CpG DNA stimulated group was treated with 1 μM K3 ODN (Endotoxin-free; Integrated DNA Technologies, Belgium), and LPS stimulated group was treated with 0.5 μg/ml LPS (Sigma, USA), as shown in Table 1.

Samples were returned to the incubator before ATR-FTIR spectroscopic analysis. Four different times points following treatment were used for measurements to evaluate the kinetics of induced effects: 10 min, 30 min, 2 h, and 6 h. When the corresponding incubation time was reached, the samples were removed from the incubator, spun at 1000 rpm, and media were discarded. Cells were washed with 15 mL of sterile PBS and spun again to ensure the complete removal of media. These steps were carried out to remove organic compounds, growth medium, or biological fluids, which can mask the IR spectrum features [58].

### 3.2. FTIR Spectroscopy

All infrared interferograms were collected using a Perkin-Elmer Spectrum One FTIR spectrometer (Perkin-Elmer Inc., Norwalk, CT, USA), with the ATR unit (Diamond-ZnSe crystal) attached. To overcome the probable atmospheric interference that can be triggered by H_2_O and CO_2_ molecules present in the air, air background was measured and subtracted automatically by the Spectrum One software (Perkin Elmer, US). The spectrum of PBS was recorded as a sample and subtracted manually. The spectra of cell samples were recorded in the 4000-650 cm^−1^ region at 25°C. Cell samples of 1 μl were dried with mild N_2_ flux for 5 min. This allows cells to settle on the crystal [30]. The best signal-to-noise ratio for interferogram collection was achieved with a 128 scan number and 2 cm^−1^ spectral resolution. ATR surface was cleaned using a water-ethanol-water sequence and then air-dried. Each group was scanned under the same conditions in three independent scans (the average constitutes a replicate n). Replicates (n=5) were used in data analysis. This protocol is available online [59].

### 3.3. Data and Chemometric Analysis

Post-processing of the spectral data—13-point Savitzky-Golay smoothing, and baseline correction at 3800-2750-1800-900 cm^−1^ points—was performed using the Spectrum One software (Perkin-Elmer). To understand the regions with significant variation, Principal Component Analysis (PCA) was performed using the spectra imported directly into the Unscrambler 11.0 (Camo, NO) software in the 3000-650 cm^−1^ region.

Peak assignment was done using the center of weight measurements and literature comparison [10]. For the side-peak band assignment, the second derivative spectrum was used. Concentration studies were carried out by integrating the curves under the peaks. For conformational alteration studies, bandwidth measurements were done manually using the 75% height of the peaks.

### 3.4. Statistical Analysis

Shapiro-Wilk and Kolmogorov-Smirnov tests were used to determine the normality of our datasets with the GraphPad Prism 8.3.1 (GraphPad, US) software. No outliers were detected nor removed. A two-way ANOVA (time point and treatment as factors) with Dunnett’s test for multiple comparisons (individual variances were computed for each comparison) were performed. P values less than or equal to 0.05 were considered as statistically significant (p ≤ 0.05 *, p ≤ 0.01 **, p ≤ 0.001 ***, and p ≤ 0.0001 ****). The results are shown as mean ± standard error of the mean (SEM).

## Supplementary Materials

Materials can be found at http://dx.doi.org/10.17632/kwnmk3fj54.1 [60]

## Author Contributions

Conceptualization, methodology, supervision, project administration, funding acquisition, F.S., M.G.; investigation, resources, software, visualization, writing—original draft preparation, E.S.; validation, data interpretation, formal analysis, data curation, writing—review and editing, S.D.; data interpretation, scientific guidance, writing—review and editing, J.S. All authors have read and agreed to the published version of the manuscript.

## Acknowledgments

The authors wish to thank Dr.Bilgi Gungor, and Dr.Soner Yildiz for cell culture preparation, Asst.Prof.Dr.Nihal Şimşek Özek and Dr.Sherif Abbas for technical help and the analysis of results.

## Abbreviations

ATR-FTIR: Attenuated Total Reflectance-Fourier Transform Infrared
LPS: Lipopolysaccharide
ODN: Oligodeoxynucleotide
PBS: Phosphate-buffered saline
HCA: Hierarchical cluster analysis
PCA: Principal component analysis

## Appendix A

**Figure A1.**
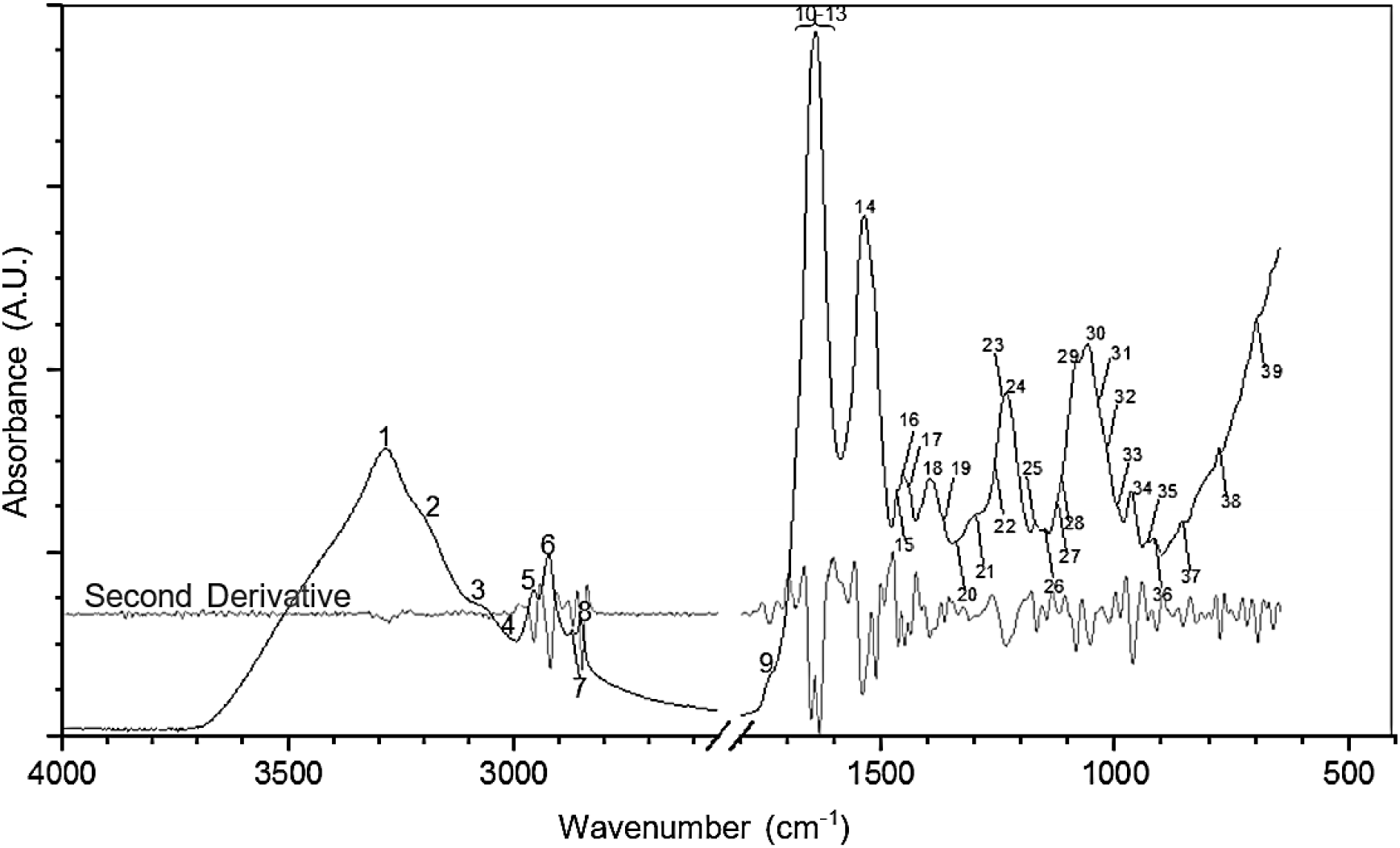
A representative FTIR spectrum of control murine macrophage cells with peak assignments in 4000-650 cm^−1^. The second derivative spectrum is obtained to analyze the shoulder peaks obscured in larger ones.

## Appendix B

**Table A1.**
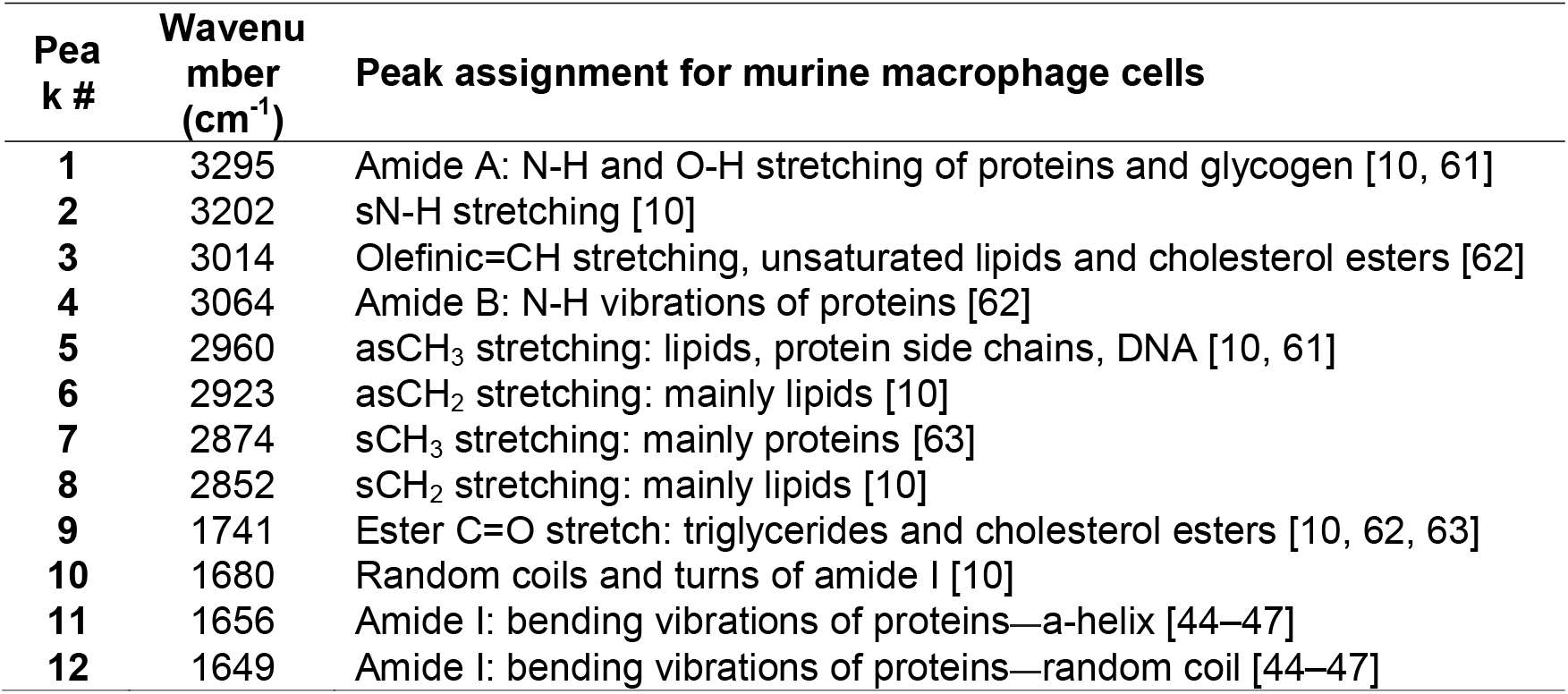
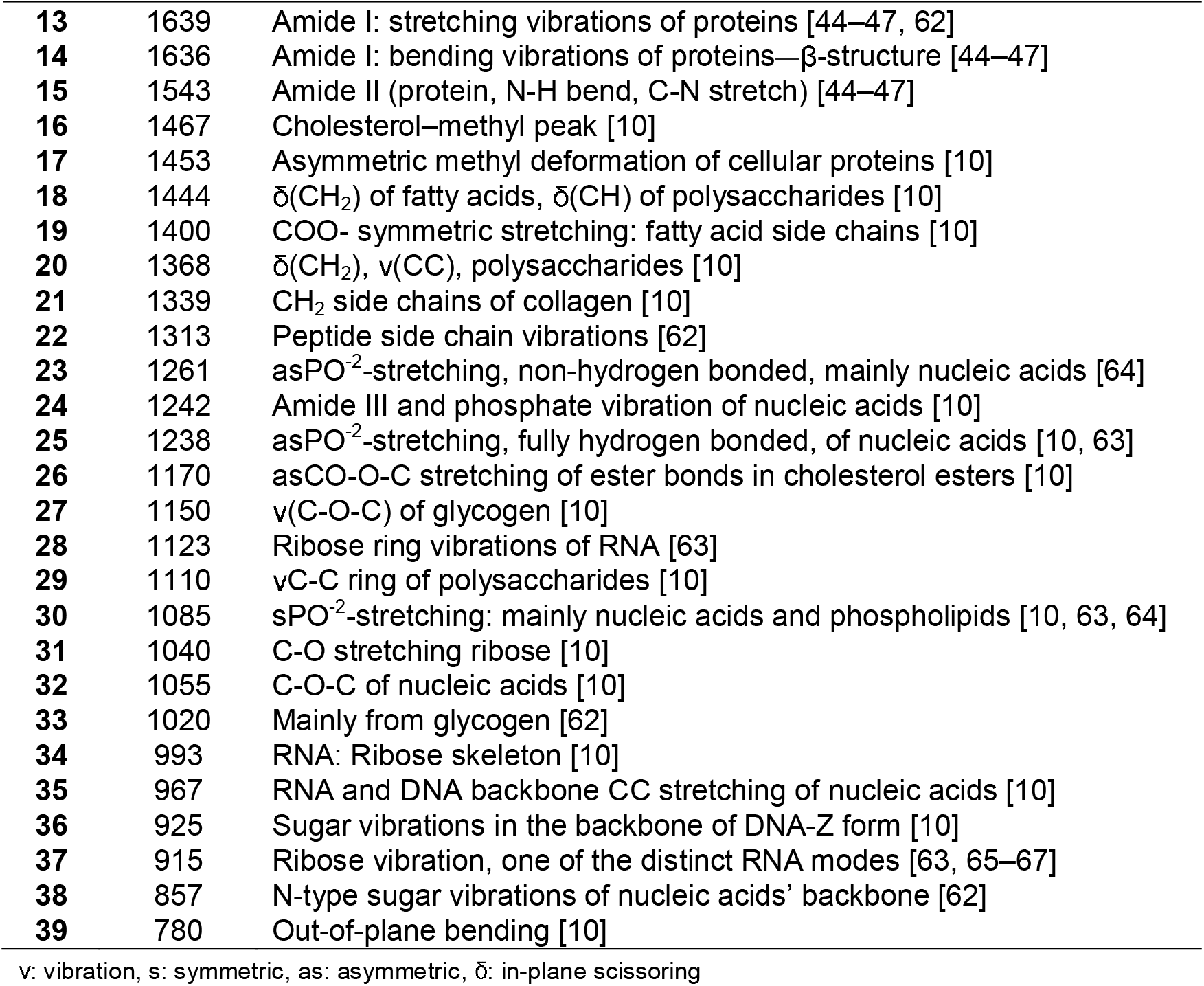
ATR-FTIR peak assignment of murine macrophage RAW 264.7 cells

| Peak # | Wavenumber ($\text{cm}^{-1}$) | Peak assignment for murine macrophage cells |
| --- | --- | --- |
| 1 | 3295 | Amide A: N-H and O-H stretching of proteins and glycogen [10, 61] |
| 2 | 3202 | sN-H stretching [10] |
| 3 | 3014 | Olefinic=CH stretching, unsaturated lipids and cholesterol esters [62] |
| 4 | 3064 | Amide B: N-H vibrations of proteins [62] |
| 5 | 2960 | asCH<sub>3</sub> stretching: lipids, protein side chains, DNA [10, 61] |
| 6 | 2923 | asCH<sub>2</sub> stretching: mainly lipids [10] |
| 7 | 2874 | sCH<sub>3</sub> stretching: mainly proteins [63] |
| 8 | 2852 | sCH<sub>2</sub> stretching: mainly lipids [10] |
| 9 | 1741 | Ester C=O stretch: triglycerides and cholesterol esters [10, 62, 63] |
| 10 | 1680 | Random coils and turns of amide I [10] |
| 11 | 1656 | Amide I: bending vibrations of proteins—α-helix [44–47] |
| 12 | 1649 | Amide I: bending vibrations of proteins—random coil [44–47] |
| 13 | 1639 | Amide I: stretching vibrations of proteins [44–47, 62] |
| 14 | 1636 | Amide I: bending vibrations of proteins—β-structure [44–47] |
| 15 | 1543 | Amide II (protein, N-H bend, C-N stretch) [44–47] |
| 16 | 1467 | Cholesterol–methyl peak [10] |
| 17 | 1453 | Asymmetric methyl deformation of cellular proteins [10] |
| 18 | 1444 | $\delta(\text{CH}_2)$ of fatty acids, $\delta(\text{CH})$ of polysaccharides [10] |
| 19 | 1400 | COO<sup>-</sup> symmetric stretching: fatty acid side chains [10] |
| 20 | 1368 | $\delta(\text{CH}_2)$, $\nu(\text{CC})$, polysaccharides [10] |
| 21 | 1339 | CH<sub>2</sub> side chains of collagen [10] |
| 22 | 1313 | Peptide side chain vibrations [62] |
| 23 | 1261 | asPO<sup>-2</sup>-stretching, non-hydrogen bonded, mainly nucleic acids [64] |
| 24 | 1242 | Amide III and phosphate vibration of nucleic acids [10] |
| 25 | 1238 | asPO<sup>-2</sup>-stretching, fully hydrogen bonded, of nucleic acids [10, 63] |
| 26 | 1170 | asCO-O-C stretching of ester bonds in cholesterol esters [10] |
| 27 | 1150 | $\nu(\text{C-O-C})$ of glycogen [10] |
| 28 | 1123 | Ribose ring vibrations of RNA [63] |
| 29 | 1110 | $\nu\text{C-C}$ ring of polysaccharides [10] |
| 30 | 1085 | sPO<sup>-2</sup>-stretching: mainly nucleic acids and phospholipids [10, 63, 64] |
| 31 | 1040 | C-O stretching ribose [10] |
| 32 | 1055 | C-O-C of nucleic acids [10] |
| 33 | 1020 | Mainly from glycogen [62] |
| 34 | 993 | RNA: Ribose skeleton [10] |
| 35 | 967 | RNA and DNA backbone CC stretching of nucleic acids [10] |
| 36 | 925 | Sugar vibrations in the backbone of DNA-Z form [10] |
| 37 | 915 | Ribose vibration, one of the distinct RNA modes [63, 65–67] |
| 38 | 857 | N-type sugar vibrations of nucleic acids' backbone [62] |
| 39 | 780 | Out-of-plane bending [10] |

v: vibration, s: symmetric, as: asymmetric, $\delta$ : in-plane scissoring

## Appendix C

**Table A2.**
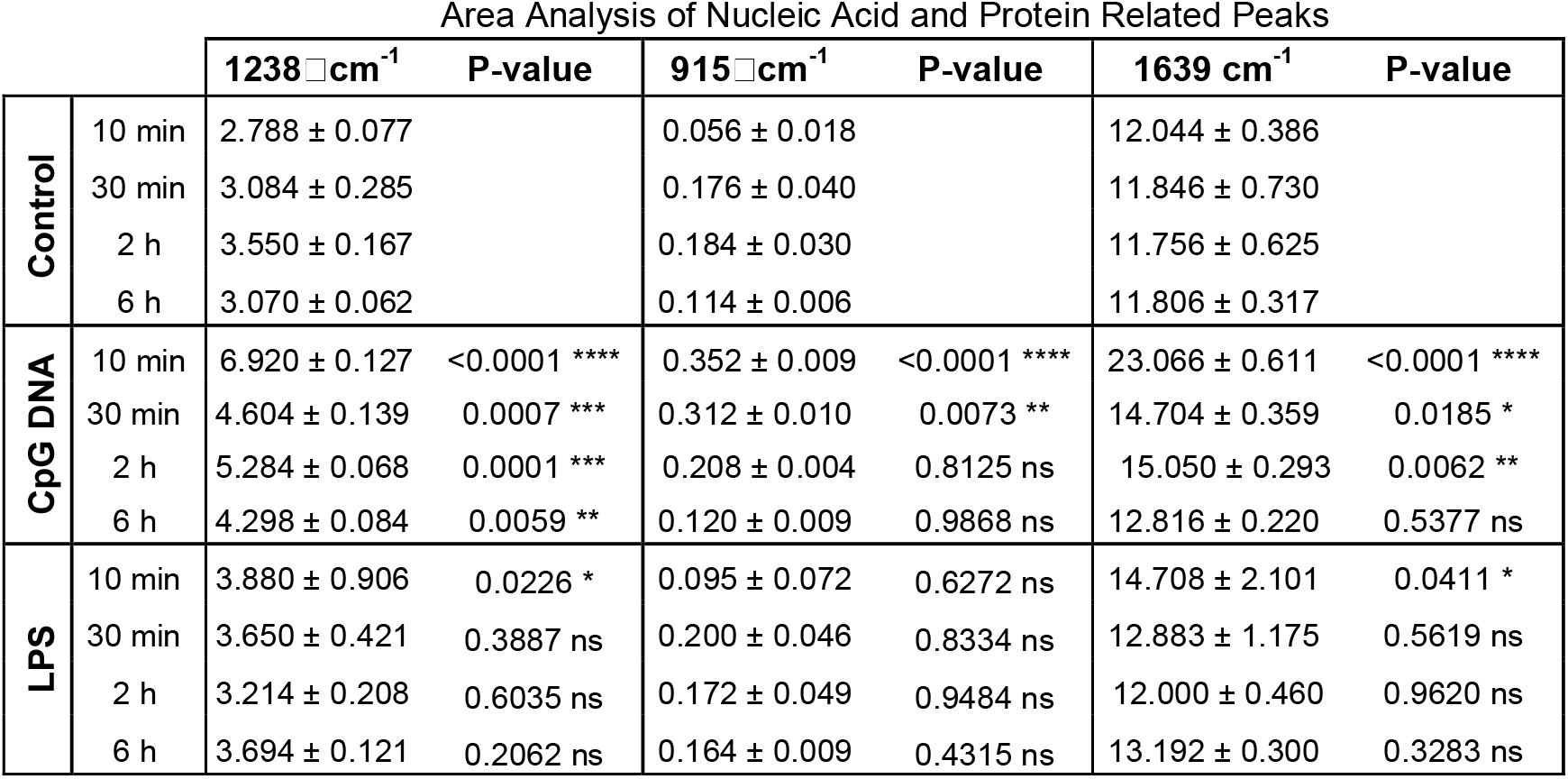
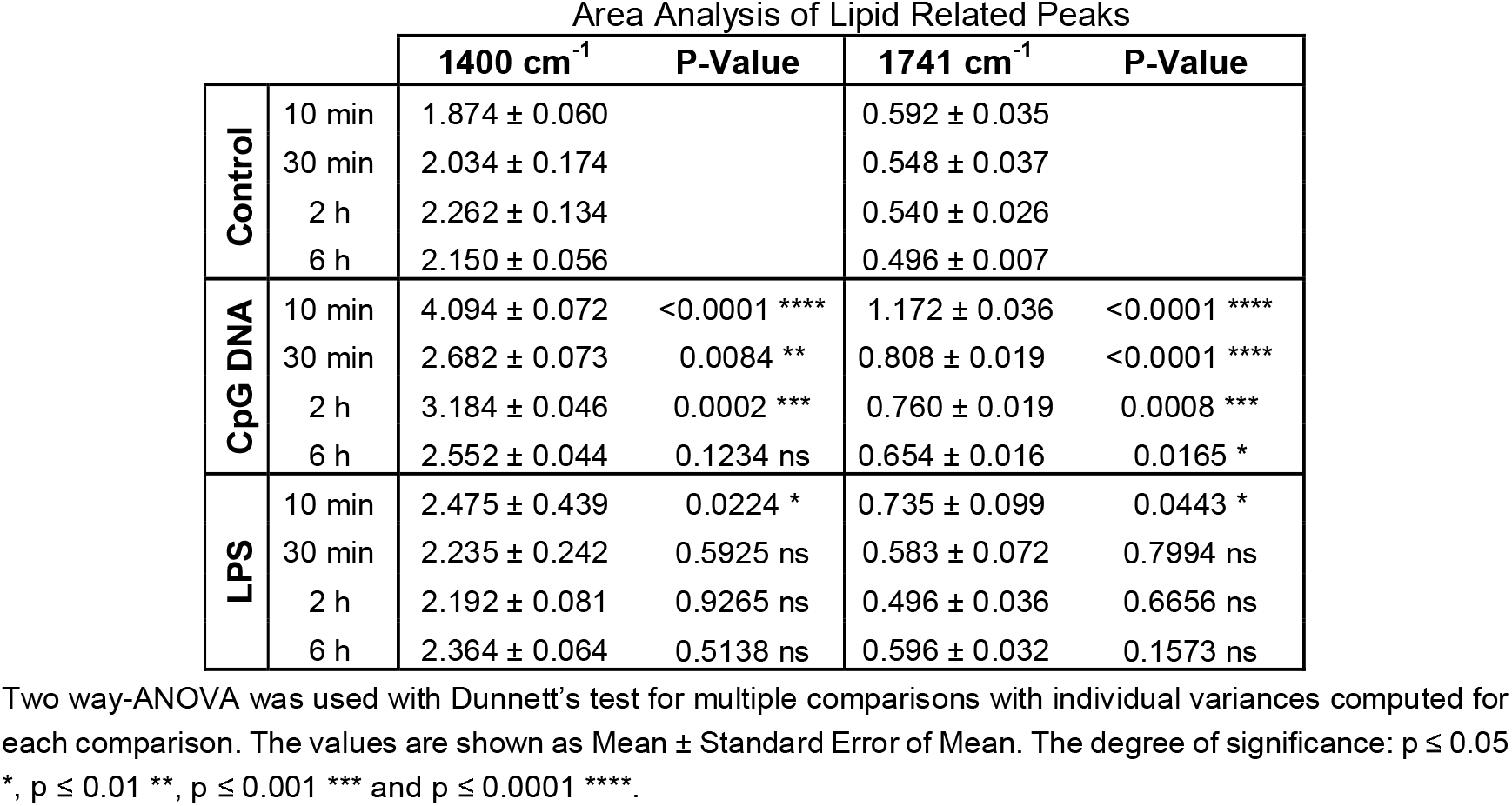
Changes in peak area values of various functional groups in control, CpG DNA, and LPS groups of RAW 264.7 cells.

| Area Analysis of Nucleic Acid and Protein Related Peaks |  |  |  |  |  |  |  |
| --- | --- | --- | --- | --- | --- | --- | --- |
| | | 1238 $\text{cm}^{-1}$ | P-value | 915 $\text{cm}^{-1}$ | P-value | 1639 $\text{cm}^{-1}$ | P-value |
| Control | 10 min | 2.788 ± 0.077 | | 0.056 ± 0.018 | | 12.044 ± 0.386 | |
| | 30 min | 3.084 ± 0.285 | | 0.176 ± 0.040 | | 11.846 ± 0.730 | |
| | 2 h | 3.550 ± 0.167 | | 0.184 ± 0.030 | | 11.756 ± 0.625 | |
| | 6 h | 3.070 ± 0.062 | | 0.114 ± 0.006 | | 11.806 ± 0.317 | |
| CpG DNA | 10 min | 6.920 ± 0.127 | <0.0001 **** | 0.352 ± 0.009 | <0.0001 **** | 23.066 ± 0.611 | <0.0001 **** |
| | 30 min | 4.604 ± 0.139 | 0.0007 *** | 0.312 ± 0.010 | 0.0073 ** | 14.704 ± 0.359 | 0.0185 * |
| | 2 h | 5.284 ± 0.068 | 0.0001 *** | 0.208 ± 0.004 | 0.8125 ns | 15.050 ± 0.293 | 0.0062 ** |
| | 6 h | 4.298 ± 0.084 | 0.0059 ** | 0.120 ± 0.009 | 0.9868 ns | 12.816 ± 0.220 | 0.5377 ns |
| LPS | 10 min | 3.880 ± 0.906 | 0.0226 * | 0.095 ± 0.072 | 0.6272 ns | 14.708 ± 2.101 | 0.0411 * |
| | 30 min | 3.650 ± 0.421 | 0.3887 ns | 0.200 ± 0.046 | 0.8334 ns | 12.883 ± 1.175 | 0.5619 ns |
| | 2 h | 3.214 ± 0.208 | 0.6035 ns | 0.172 ± 0.049 | 0.9484 ns | 12.000 ± 0.460 | 0.9620 ns |
| | 6 h | 3.694 ± 0.121 | 0.2062 ns | 0.164 ± 0.009 | 0.4315 ns | 13.192 ± 0.300 | 0.3283 ns |

| Area Analysis of Lipid Related Peaks |  |  |  |  |  |
| --- | --- | --- | --- | --- | --- |
|  |  | 1400 cm<sup>-1</sup> | P-Value | 1741 cm<sup>-1</sup> | P-Value |
| Control | 10 min | 1.874 ± 0.060 |  | 0.592 ± 0.035 |  |
|  | 30 min | 2.034 ± 0.174 |  | 0.548 ± 0.037 |  |
|  | 2 h | 2.262 ± 0.134 |  | 0.540 ± 0.026 |  |
|  | 6 h | 2.150 ± 0.056 |  | 0.496 ± 0.007 |  |
| CpG DNA | 10 min | 4.094 ± 0.072 | <0.0001 **** | 1.172 ± 0.036 | <0.0001 **** |
|  | 30 min | 2.682 ± 0.073 | 0.0084 ** | 0.808 ± 0.019 | <0.0001 **** |
|  | 2 h | 3.184 ± 0.046 | 0.0002 *** | 0.760 ± 0.019 | 0.0008 *** |
|  | 6 h | 2.552 ± 0.044 | 0.1234 ns | 0.654 ± 0.016 | 0.0165 * |
| LPS | 10 min | 2.475 ± 0.439 | 0.0224 * | 0.735 ± 0.099 | 0.0443 * |
|  | 30 min | 2.235 ± 0.242 | 0.5925 ns | 0.583 ± 0.072 | 0.7994 ns |
|  | 2 h | 2.192 ± 0.081 | 0.9265 ns | 0.496 ± 0.036 | 0.6656 ns |
|  | 6 h | 2.364 ± 0.064 | 0.5138 ns | 0.596 ± 0.032 | 0.1573 ns |

Two way-ANOVA was used with Dunnett's test for multiple comparisons with individual variances computed for each comparison. The values are shown as Mean ± Standard Error of Mean. The degree of significance: $p \leq 0.05$ \*, $p \leq 0.01$ \*\*, $p \leq 0.001$ \*\*\* and $p \leq 0.0001$ \*\*\*\*.

